# Representations of temporal community structure in hippocampus and precuneus predict inductive reasoning decisions

**DOI:** 10.1101/2021.10.12.462707

**Authors:** Athula Pudhiyidath, Neal W Morton, Rodrigo Viveros Duran, Anna C. Schapiro, Ida Momennejad, Demitrius M. Hinojosa-Rowland, Robert J. Molitor, Alison R. Preston

## Abstract

Our understanding of the world is shaped by inferences about underlying structure. For example, at the gym, you might notice that the same people tend to arrive around the same time and infer that they are friends that work out together. Consistent with this idea, after participants are presented with a temporal sequence of objects that follows an underlying community structure, they are biased to infer that objects from the same community share the same properties. Here, we used fMRI to measure neural representations of objects after temporal community structure learning and examine how these representations support inference about object relationships. We found that community structure learning affected inferred object similarity: when asked to spatially group items based on their experience, participants tended to group together objects from the same community. Neural representations in perirhinal cortex predicted individual differences in object grouping, suggesting that high-level object representations are affected by temporal community learning. Furthermore, participants were biased to infer that objects from the same community would share the same properties. Using computational modeling of temporal learning and inference decisions, we found that inductive reasoning is influenced by both detailed knowledge of temporal statistics and abstract knowledge of the temporal communities. The fidelity of temporal community representations in hippocampus and precuneus predicted the degree to which temporal community membership biased reasoning decisions. Our results suggest that temporal knowledge is represented at multiple levels of abstraction, and that perirhinal cortex, hippocampus, and precuneus may support inference based on this knowledge.

## Introduction

The world is richly structured in many ways that may help guide decision making; however, this structure is not always directly observable and must often be inferred based on limited data. The sequence in which events occur may be an important factor in providing clues about underlying structure (Gershman et al., 2017; Schapiro et al., 2013; Varga et al., n.d.). For example, when at the gym, you might notice that a group of people tends to enter near the same time. Based on this observation, you might infer that they are friends that work out together. This inference of membership within a friend group could then be used to support further reasoning; for example, if you learn that one member of the group is a student at a local university, you might then assume that other members of the group also go to that university. In this way, knowledge about temporal statistics may influence reasoning about non-temporal structure in the world.

Recent studies have used a temporal community structure paradigm to examine learning of latent structure through statistical learning (Schapiro et al., 2013, 2016) and determine how this learning may affect reasoning (Pudhiyidath et al., 2019). In the temporal community structure paradigm, participants are presented with sequences of items that are generated according to an underlying network structure that defines connections between items (Fig. 1a). After presentation of each item, any other connected item may be presented next with equal probability (Fig. 1b). Items within a community are interconnected, and there are only sparse connections between communities; as a result, items in the same community tend to appear nearby in time to one another. After observing item sequences, participants are more likely to report perceived event boundaries at transitions between communities, indicating that they have learned about the latent community structure of the items (Schapiro et al., 2013). Learning of temporal community structure also biases subsequent inductive reasoning about items (Pudhiyidath et al., 2019). After presentation of item sequences generated based on a temporal community structure, participants were told a property of a cue item (e.g., that it likes the desert) and asked to decide which of two other items shares that property. Adults were more biased than children to select an item that shared the same temporal community with the cue item. This result suggests that, in adults, temporal knowledge about item communities influenced reasoning about non-temporal properties of the items (Pudhiyidath et al., 2019). However, the specific computational and neural mechanisms that support generalization of temporal knowledge to make decisions in new contexts remain unclear. Here, we used fMRI and computational modeling to investigate how temporal knowledge is represented in the brain and how this knowledge affects inference decisions.

**Figure 1.**
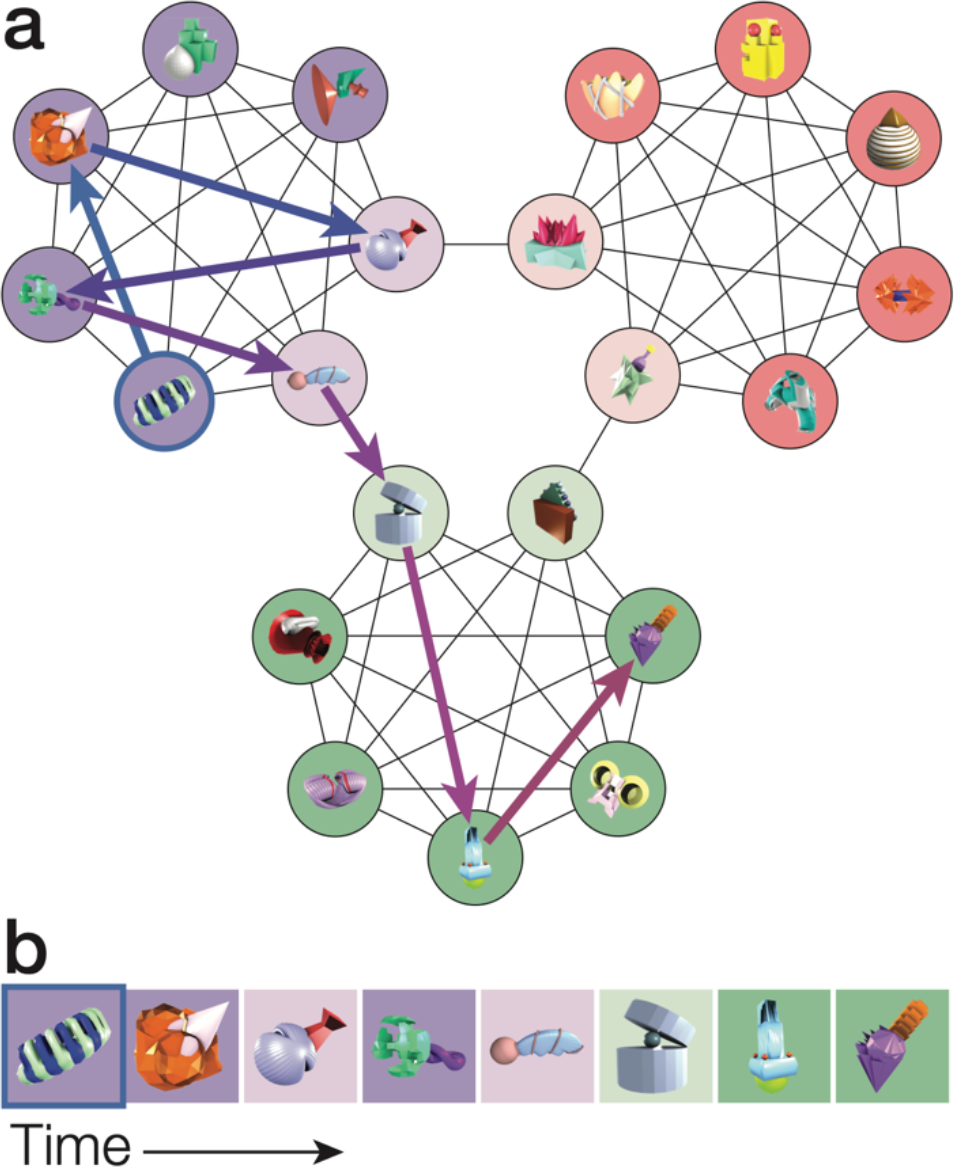
**(a)** Temporal community structure with novel 3D objects assigned to its 21 nodes. The connections between nodes indicate which objects may follow one another in succession in a structured sequence. Nodes are divided into three distinct communities (purple, red, green). Darker circles indicate central nodes, i.e., objects that are connected to all other objects in the same community. Lighter circles indicate boundary nodes, i.e., objects that are connected to the five central nodes and another boundary object in a different community. The arrows show an example sequence through the network, starting with the outlined object; arrow color corresponds to position within the sequence. **(b)** A structured sequence of objects corresponding to the path illustrated in (a). For illustrative purposes, background color corresponds to the node colors in (a). In the actual experiment, objects were presented on a white background.

Computational models of associative learning and reinforcement learning propose that items that are presented nearby in time to one another become associated in memory (Dayan, 1993; Howard & Kahana, 2002; Raaijmakers & Shiffrin, 1980). Gradual learning of temporal associations may then allow participants to form predictions of what items will follow any given presented item (Gershman et al., 2012; Schapiro et al., 2013, 2017) to support efficient decision making (Dayan, 1993; Momennejad, 2020; Momennejad et al., 2017). Successor representation (SR) models propose that predictions about upcoming states or items are stored in an SR that reflects not only one-step transitions, but also longer-range transitions (Dayan, 1993). For any given item, the SR stores the number of expected future visits to every other item. Expected visits are discounted based on the expected delay before visiting them, causing the SR to emphasize items that will be visited relatively soon. When an SR model was trained on the temporal community structure task, it learned predictive representations that reflected the community structure, associating items in the same community more strongly than items in different communities (Stachenfeld et al., 2017). This result suggests that SR learning might be able to account for influences of community structure on event segmentation (Schapiro et al., 2013) and inference (Momennejad, 2020; Pudhiyidath et al., 2019). However, while the SR is sensitive to the community structure, it also represents other details of the temporal statistics of the paradigm, such as the connections between communities (Stachenfeld et al., 2017). The SR has been theorized to support formation of compressed representations that are more abstract (Momennejad, 2020; Stachenfeld et al., 2017). Therefore, SR learning might support reasoning based on community structure either directly through knowledge of temporal statistics, or indirectly by supporting learning of abstract structure (here, the community membership of each item).

Learning of temporal statistics is thought to be facilitated by the hippocampus (Schapiro et al., 2014; Schlichting et al., 2017), which has been proposed to encode SRs that reflect predictions about upcoming states (Brunec et al., 2018; Stachenfeld et al., 2017). Hippocampal activation patterns in response to objects reflect the co-occurrence statistics of those objects: Objects that frequently followed one another during learning come to have more similar activation patterns after learning (Schapiro et al., 2012). Furthermore, recent work suggests that the hippocampus supports memory performance in statistical learning tasks. Developmental changes in anterior hippocampus volume predict individual differences in statistical learning performance across development (Schlichting et al., 2017), and statistical learning is disrupted in patients with hippocampal damage (Covington et al., 2018; Schapiro et al., 2014). After exposure to sequences of objects generated based on temporal community structure, patterns of activation in the hippocampus reflect community membership, with greater similarity among items in the same community compared to items in different communities (Schapiro et al., 2016). However, the behavioral relevance of these community representations remains unclear, as the relationship between hippocampal community structure representations and reasoning behavior has not yet been investigated.

The prefrontal cortex (PFC) has also been found to represent community structure after learning. A previous study found that the inferior frontal gyrus (IFG) increased in activation as multiple items from the same community were presented in succession, suggesting that it is sensitive to community structure (Schapiro et al., 2013). IFG activation patterns also reflected community membership, with greater observed similarity among object presentations from the same community compared to objects in different communities (Schapiro et al., 2013).

However, these results were observed during presentation of ordered sequences that visited all objects in the same community in succession, and IFG activation was sensitive to the position within these sequences. Therefore, it remains unclear whether IFG represents the community membership of individual objects or instead represents a context-dependent activation pattern reflecting the current community. There is also evidence that medial prefrontal cortex (mPFC) is sensitive to community structure. While Schapiro and colleagues did not find evidence that mPFC activation patterns are more similar for items from the same community compared to items from different communities, mPFC activation was increased during a sequence of object presentations from the same community compared to boundaries between communities (Schapiro et al., 2013). Furthermore, in studies on prospective memory, the mPFC has been found to represent intended upcoming tasks (Momennejad & Haynes, 2012, 2013), raising the possibility that it might also represent predicted upcoming items within a given community. The mPFC is also functionally and structurally connected to hippocampus and has been proposed to form low-dimensional representations of task structure (Mack et al., 2020; Morton et al., 2020; Morton & Preston, 2021). These findings raise the possibility that mPFC may form an abstract representation of task structure that reflects object community membership.

To examine how neural representations of temporal knowledge relate to reasoning behavior, we collected fMRI data in a temporal community structure paradigm. We first presented participants with sequences of objects drawn from an underlying temporal community structure (Fig. 1). We then used fMRI to measure neural representations of the objects. While prior work measured object representations during predictable sequences that followed the community structure of the task (Schapiro et al., 2013, 2016), here we measured patterns of activation elicited by the objects presented in random order (Fig. 2a). This strategy allowed us to measure the representation of each object separately without any influence of sequence-based predictions, to determine whether representations of individual objects reflect learning of the community structure. We examined object pattern similarity in hippocampus, IFG, and mPFC to test whether objects in the same community were represented more similarly to each other compared to objects in different communities.

**Figure 2.**
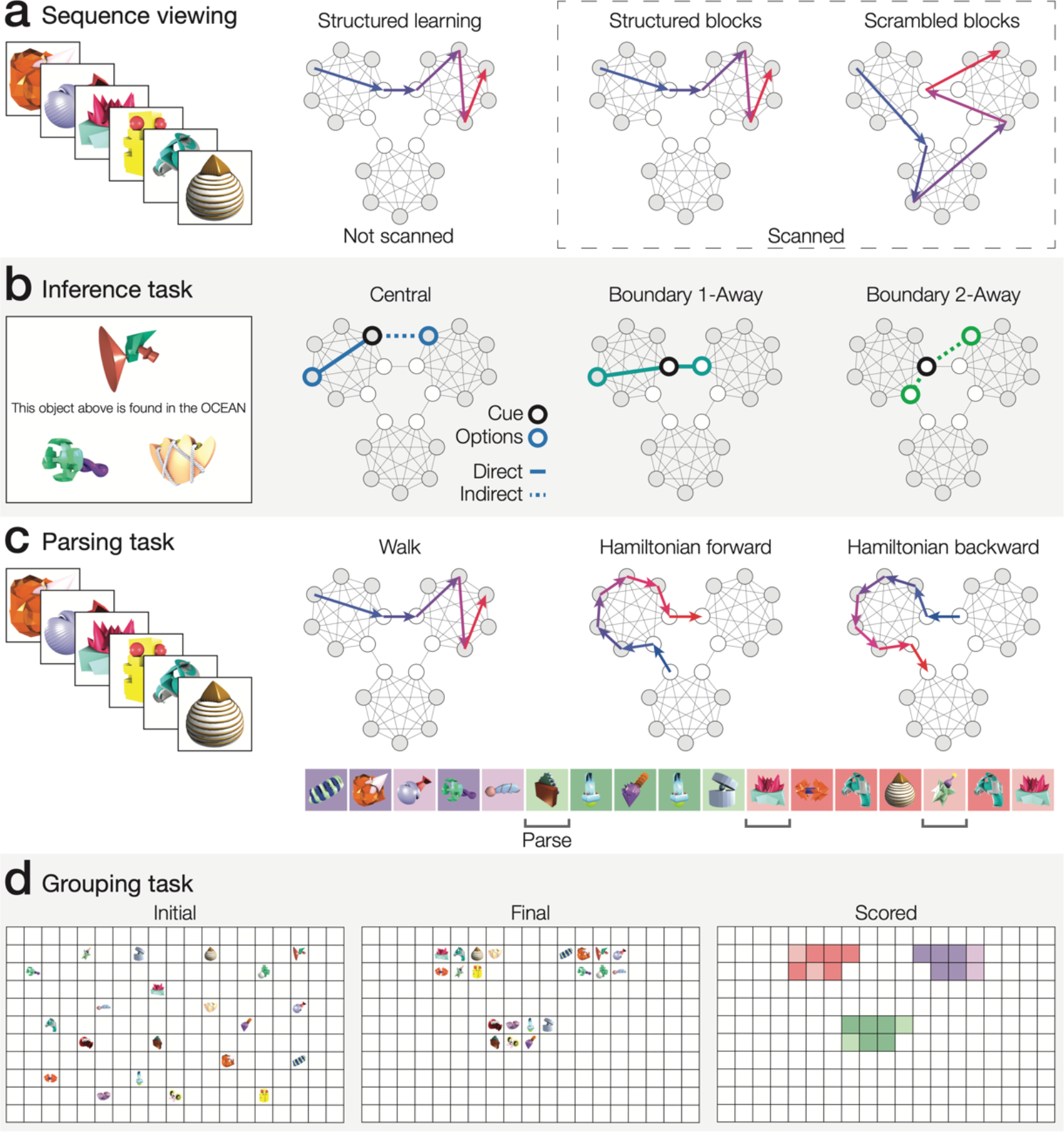
**(a)** Participants first viewed structured sequences of objects derived from the temporal community structure graph. Next, participants were scanned during presentations of additional structured sequence blocks, which were intermixed with scrambled sequence blocks in which objects were presented in a random order that did not follow the temporal community structure. **(b)** Outside of the scanner, participants performed an inference task to measure how temporal structure learning affected reasoning about object properties. Participants were told that a cue object was found in one of three environments (ocean, desert, or forest) and asked to choose which of two objects (shown below the cued object) could also be found there. For each trial, one of the object choices shared the same temporal community as the cue, and the other object choice did not. There were three types of trials (central, boundary 1-away, and boundary 2-away) which varied in the degree to which the choice objects were directly or indirectly connected with the cue object in the structure graph. **(c)** The parsing task measured subjective perception of event boundaries. Participants were presented with object sequences that were generated either through a structured walk through the graph or a Hamiltonian walk in which the nodes were visited one-by-one in turn. Participants were told to press a button (i.e., indicate an event boundary) whenever they felt a subjective shift in the sequence. In the example sequence, the object’s background color (not shown to participants) indicates its community; dark squares indicate central nodes and light squares indicate boundary nodes. **(d)** The grouping task measured the learned similarity of the objects and examined whether temporal knowledge generalized to a spatial grouping task. Participants were shown a grid with the 21 objects they had seen in the task randomly placed on it and were asked to group the objects based on their experience.

We next examined whether neural representations of community structure predicted decisions that require making inferences about object similarity and object properties.

Participants were asked to group objects within a spatial grid based on their experience with the objects, allowing us to examine whether temporal community structure learning affected the inferred similarity of the objects (Fig. 2d). We tested whether participants tended to group together items from the same temporal community and examined whether this tendency was predicted by neural representations of temporal community structure. We also examined whether learning about the temporal associations of the objects affected inductive reasoning about object properties (Fig. 2b).

During the reasoning task, participants were told a property of a cue object and asked to select which of two candidate objects shared that property. Each choice was between an object in the same community as the cue and an object from a different community. Critically, there was no “correct” answer and participants did not receive feedback; this design allowed us to examine how temporal knowledge biases reasoning in new situations. Different trials were designed to examine relationships between objects that were either directly associated during learning (i.e., directly connected in the community structure) or only indirectly associated (i.e., members of the same temporal community, but not directly connected to one another in the structure and therefore not seen together during learning). Contrasting performance on these different trials allowed us to determine whether reasoning was related only to the transition statistics that participants learned directly, or whether their reasoning was also biased by inferred associations between objects that were in the same community but were not connected directly.

To examine the influence of different forms of temporal knowledge on reasoning behavior, we compared different models of temporal knowledge based on either SR learning or abstract knowledge about latent community structure. Finally, we tested whether neural representations of temporal community structure in hippocampus, IFG, or mPFC predicted reasoning performance. Overall, our approach allowed us to determine how learning about temporal statistics influences inference-based decision making in new contexts.

## Methods

### Participants

Thirty-nine right-handed volunteers participated in the experiment (20 women; ages 18 to 33 years; M = 24.59, SD = 4.62). Consent was obtained in accordance with an experimental protocol approved by the Institutional Review Board at the University of Texas at Austin.

Participants received monetary compensation for their involvement in the study. Data from two participants were excluded for excessive movement. Data from the remaining 37 participants were included in all subsequent analyses (18 women; ages 18 to 33 years; M = 24.68, SD = 4.80).

### Stimuli

The stimuli consisted of images of 21 novel, computer generated 3D objects, selected from a larger set of previously developed object images (Hsu et al., 2014; Schlichting et al., 2015). The images of the objects were 400 pixels by 400 pixels. Object images were chosen to minimize visual similarity (e.g., color, shape) between objects. At the outset of the experiment, participants completed a familiarization task, wherein they learned the canonical orientation of each object. On each trial of the familiarization task, an object was presented for 3 s.

Participants were instructed to pay close attention to the features and orientation of each object because they would later be asked to make judgments about object orientation. Between trials, a fixation cross was shown during an inter-stimulus interval (ISI) of 0.5 s. The 21 objects were randomly ordered, and the random sequence was presented twice.

### Temporal community structure

To investigate how temporal statistical learning affects reasoning, we presented participants with object sequences generated based on a temporal community structure (Schapiro et al., 2013). Each object was assigned to a node of a 21-node community structure graph (Fig. 1a) that specified the connections among the objects (Schapiro et al., 2013). The assignment of objects to nodes was randomized for each participant. In the temporal community structure, there were three distinct communities of objects that were connected to one another. Within each of the three communities, there were five central nodes and two boundary nodes. Each node was connected to six other nodes. The central nodes were connected to all the other objects in that community. The boundary nodes were each connected to all of the central nodes in that community and to a boundary node from a different community. The two boundary nodes in each community were not connected to one another.

The community structure was used to generate structured object sequences through random walks following the connections within the graph (Fig. 1b). During structured object sequences, after presentation of an object, the following object could be any of the six connected objects with equal probability. Central node objects were always followed by another object in the same community, while boundary objects had a 5/6 probability of transitioning to another object in its own community and a 1/6 probability of transitioning to another, connected community. The structured object sequences thus provided an opportunity to learn about both the transition probabilities among individual pairs of objects and the overall latent temporal community structure.

### Learning the temporal community structure

After the object familiarization task, participants viewed structured object sequences generated from walks through the community structure (Fig. 1). The temporal structure learning task was divided into two parts. The first part, which occurred in a quiet testing room, provided participants with an opportunity to implicitly learn the underlying temporal community structure. The second part occurred during fMRI data acquisition, allowing us to measure how neural representations of the objects were affected by community structure learning.

### Initial structure learning

During initial structure learning, each participant viewed a sequence of the novel 3D objects. Each object was presented for 1.5 s with no ISI. Unbeknownst to the participants, the object sequence was generated based on a temporal community structure (Fig. 1). For each participant, a structured sequence of 1575 objects was randomly generated following the graph structure; this sequence was then divided into five runs with 315 object presentations each.

Across the five runs, each object was presented 75 times on average (standard deviation: M=17.39, SE=1.15), and there were 525 object presentations from each community on average (standard deviation: M=129.94, SE=10.42). Participants were given a chance to take a short break after each run. While viewing the sequence, participants completed an object rotation detection task. Participants made a response to every object presented, pressing one button if the object was in the canonical orientation they had learned during the familiarization phase or another button if it was in the incorrect, rotated orientation. Rotated object images were rotated 90° counterclockwise from the canonical image. Unbeknownst to participants, objects were rotated when the current object was the same as the object presented two steps back. At the end of each run, participants were given feedback about their accuracy in the rotation detection task before they moved onto the next run. To confirm that participants were consistently attending to the object presentations, performance on the rotation detection task for each run was assessed using *d*′. For each participant and run, we tested whether performance was above chance using a permutation test. On each of 10000 iterations, the responses were permuted and *d*′ was calculated based on the permuted responses. We then calculated a *d*′-value based on the permutation distribution to test whether performance was above chance on each run. To assess whether fatigue affected performance during later runs, we also tested whether *d*′ varied across runs using a one-way repeated measures ANOVA.

### Scanned structure learning

Next, participants continued viewing object sequences and performing the rotation detection task while undergoing fMRI scanning, allowing us to measure the neural representations of the objects after exposure to the temporal community structure. Participants completed six functional scanning runs, each lasting approximately 10 minutes. Unbeknownst to participants, each run was divided into structured and scrambled blocks. Structured blocks were generated based on the temporal community structure graph, as in the initial community structure learning phase. In contrast, scrambled blocks did not follow the temporal community structure (Fig. 2a).

During scrambled blocks, each of the 21 objects was presented in random order. These scrambled blocks allowed us to measure the neural representation of each object while controlling for effects of temporal autocorrelation (Purdon & Weisskoff, 1998), which were confounded with community structure during the structured blocks. Furthermore, prior work found that neural activation varied when multiple objects from the same temporal community were presented in succession (Schapiro et al., 2013). Here, we used the scrambled blocks to examine individual object representations in the absence of systematic order effects and determine how they have been shaped by temporal community learning.

Each of the six runs contained seven blocks of 21 objects each, including five structured blocks and two scrambled blocks, for a total of 147 object presentations per run. Across the six runs, each object was presented 42 times on average (standard deviation: M=9.32, SE=0.58), and there were 294 object presentations from each community on average (standard deviation: M=65.40, SE=5.54). Each object presentation lasted for 1 s, during which participants pressed one button to indicate that the object was in the canonical orientation or another button to indicate that the object was rotated from its canonical orientation. Participants were not given feedback about their performance during the scanned phase. As with the initial structure learning task, we assessed performance on the orientation task by calculating *d*′. After each trial, a fixation cross was presented during a 1, 3, or 5 s ISI. There were an equal number of ISIs of each length in each run, with the order of ISI lengths randomized. Each run began and ended with 4 s of fixation. Within each run, the scrambled blocks occurred as either the third and sixth blocks (schedule 1), or as the second and fifth blocks (schedule 2); the block order started with schedule 1 and alternated across runs. The start of each structured block followed the temporal community graph, based on the object presented previously. All participants were given a short break (approximately 10 minutes) after completing the first three runs in the scanner to avoid fatigue, after which they returned to the scanner to complete the final three runs.

### Assessing knowledge of temporal community structure

After the scanned structure learning phase, a series of behavioral tasks assessed knowledge of the temporal statistics of the objects and examined how temporal knowledge affects decisions that rely on inference. We measured temporal knowledge using a parsing task that characterized whether temporal community structure affected event segmentation. We further examined whether temporal knowledge was generalized to a spatial grouping task, which assessed the subjective similarity of the objects. Finally, we tested whether temporal statistical learning affected reasoning using an inductive inference task. To avoid influences of the other tasks on inductive inference, the inductive inference task was completed first, followed by the parsing task and the spatial grouping task.

### Parsing task

The parsing task allowed us to assess whether temporal community structure learning affected perception of event boundaries. In this task, participants were shown object sequences and told to press a button (i.e., parse the sequence) whenever they believed that there was a “shift” in the object sequence (Fig. 2c) (Schapiro et al., 2013). Participants were told to use their intuition and to refer to their experiences thus far in the experiment to determine where shifts might be occurring. Based on prior work (Schapiro et al., 2013), we predicted that participants would be more likely to make a parse response after transitions between communities.

During the parsing task, participants viewed object sequences, which were divided into three runs of 252 objects. Each object was shown on the screen for 1.5 s, with no ISI between objects. Each run contained 12 blocks with 21 objects each. Each block was either a random walk through the temporal graph structure (i.e., a structured sequence like those encountered during structure learning) or a Hamiltonian walk (Fig. 2c). In Hamiltonian walks, participants saw each object in the graph in a sequential order that followed the outer edges of the graph. Within each run, blocks alternated between random walks and Hamiltonian walks, starting with a random walk. Each Hamiltonian walk was randomly selected to follow either a predetermined forward order or a backward version of the same walk (Schapiro et al., 2013). These Hamiltonian paths were included to ensure that parsing responses could not be solely explained by changes in the temporal statistics of object presentations, as each object was only presented once in each block. In contrast, during random walks, objects in the same community are more likely to repeat in relatively quick succession, while objects in a new community likely have not been observed recently. As a result, parsing responses might occur more frequently after community transitions because of object novelty rather than knowledge about temporal structure.

To measure the influence of temporal community structure on event segmentation, we examined the probability of making a parse response after a transition from one community to another. Following prior work (Schapiro et al., 2013), we counted a community transition only after there had been at least four objects presented from the same community in succession, in case participants were disinclined to make parse responses in close succession to one another. We compared the probability of parsing on these community transition trials (i.e., when there were at least four trials with objects in the same community, followed by presentation of an object in a different community) compared to the probability of parsing on other trials. We calculated this difference in parsing probability separately for the structured and Hamiltonian blocks. The difference in parsing probability between community transition trials and other trials served as a measure of knowledge of the temporal community structure.

### Grouping task

In the grouping task, participants were asked to group objects within a spatial grid based on their experiences in the study. The grouping task involved inferring the relative strength of the relationships among all the objects, based on limited information obtained during the temporal structure learning task. This task thus allowed us to assess whether temporal knowledge influenced inference behavior in a spatial task. Participants were shown a 19x11 rectangular grid on a computer screen, with the 21 objects uniformly and randomly dispersed across its coordinates (Fig. 2d). Participants were instructed to use the computer mouse to group the objects on the grid. They were asked to group the objects based on their intuition and what they had witnessed thus far in the experiment and were instructed to not use the physical similarity of the objects to guide their responses. To group the objects, participants clicked on an object and then clicked on the space on the grid where they wanted to move it. Participants were given as long as they needed to complete the task.

Using the final positions of the objects, we calculated the Euclidean distance between objects in the same community and objects in different communities. We hypothesized that the spatial distances between objects would reflect the temporal community structure, with shorter distances between objects in the same community. We found that there was substantial individual variability in how participants used the grid to separate groups. For example, some participants placed groups in different corners of the grid, while others placed all objects in a small area, using less than a third of the grid. Therefore, we focused mainly on the distances among objects in the same community, which are less sensitive to individual differences in the strategy used to separate groups.

### Inductive inference task

During the inductive inference task, we tested whether learning of the temporal relationships between objects influenced reasoning about properties of the objects. Specifically, participants were told the habitat where a cue object is found (ocean, desert, or forest) and then were asked to infer which of two different choice objects shared the same habitat as the cue (Fig. 2b). On each inference trial, the cue object appeared at the top of the screen with a statement about its environment (e.g., “This object above is found in the FOREST.”), with the two choice objects shown at the bottom of the display. Unbeknownst to the participants, the stated habitat of each cue object corresponded to the community it belonged to in the graph structure. On each inference trial, one of the choices was another object from the same temporal community as the cued object, while the other choice was an object from a different community to the cued object. This task structure allowed us to examine whether participants were biased to infer shared properties among objects from the same community. On each trial, participants were given 8 s to select one of the two choice objects with a button press. After the participant responded, a fixation cross was presented for 0.5 s, followed by the next trial.

Each participant completed 42 inference trials. Each of the 21 objects was used as the cue object on two trials. Inference trials were divided into three different trial types: central, boundary 1-away, and boundary 2-away (Fig. 2b). These trial types were designed to examine both the influence of knowledge about individual transitions between object pairs (i.e., the frequency with which a pair of objects was adjacent in the sequence) and the influence of abstract knowledge about the temporal community structure (i.e., whether two objects were in the same community or different communities). On each of the 30 central inference trials, each of the objects was a central node within the temporal graph structure. Participants were given a choice between (1) a central object in the same community as the cue and (2) a central object from a different community. The central trials therefore involved choosing between an object that was directly connected to the cue and in the same community, and an object that was only indirectly connected and in a different community.

In contrast, the boundary trials controlled for direct associations between objects while examining whether participants were influenced by knowledge of the community structure. In the six boundary 1-away trials, the cue object was a boundary node in the community graph, and the choice objects were both directly connected to the cue object in the graph (i.e., they were 1 connection away from the cue object). Participants chose between (1) a central object from the same community as the cue and (2) a boundary object from a different community. Finally, there were six boundary 2-away trials, in which the cue object was a boundary node and the choice objects were only indirectly connected to the cue object in the graph, with a minimum path length of two connecting the cue object to each of the choice objects. Participants chose between (1) the other boundary object in the same community as the cue and (2) a central object in a different community. Notably, on boundary 2-away trials, neither choice object was directly associated with the cue item; as a result, any temporal bias observed on these trials could not be based on direct associations.

For each trial type, we examined whether participants were biased by the temporal community structure to infer that objects in the same community were from the same environment. We measured this bias using a temporal bias score defined as the proportion of trials for which the participant selected the same-community object minus the proportion of trials for which the participant selected the different-community object. A bias score of 1 indicates maximum bias toward inferring shared properties among same-community objects, while a bias score of 0 indicates no temporal bias. Given the relatively small number of inference trials, we did not anticipate that inference decisions would be affected by learning of the environment associated with each cue object. Nevertheless, to confirm that inference decisions reflected prior structure learning rather than learning during the inference phase, we tested whether temporal bias varied with time during the inference task. If learning did occur during the inference task, then performance would increase during later trials. We divided the 42 inference trials into six bins of seven trials each, and used a one-way repreated-measures ANOVA to test whether temporal bias (collapsed over trial types) varied with trial bin.

### Modeling of temporal bias in the inference task

To better understand how participants make inductive inferences about shared object properties, we simulated inference task performance using different competing models. The inductive inference task examined the effects of direct associations and indirect associations, allowing us to contrast different forms of temporal knowledge that might bias reasoning decisions. Each model assumed that inference of shared object properties was proportional to the strength of association of the objects in memory. We considered two models of association strength: successor representation (SR) and community membership. The SR stores the temporally discounted expected frequency with which each object would be visited, conditional on a given object having been presented (Dayan, 1993; Momennejad, 2020). The SR was initialized as an *n* x *n* matrix of zeros, where *n* is the number of objects. The SR was then updated using temporal difference learning during presentation of object sequences during the structure learning task, according to

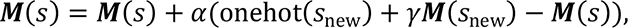

where *M*(*s*) is the SR for state *s*, *α* is a learning rate parameter (0 < *α* < 1), *γ* is a discount parameter (0 < *γ* < 1) that determines the predictive horizon of the SR, and onehot(*s*_new_) is a vector of all zeros with a 1 for successor state *s*_new_. We simulated learning through the five runs of the initial structure learning and the six runs of the scanned structure learning tasks, updating the SR based on the object sequences presented in each run. Note that, in the scanned structure learning tasks, the scrambled blocks may cause some unlearning of the community structure; this unlearning was accounted for in the model. To examine how the model parameters affected the SR, we simulated SR learning using a range of values for the *α* and *γ*parameters, based on the object sequences presented to a sample participant (Fig. 3).

**Figure 3.**
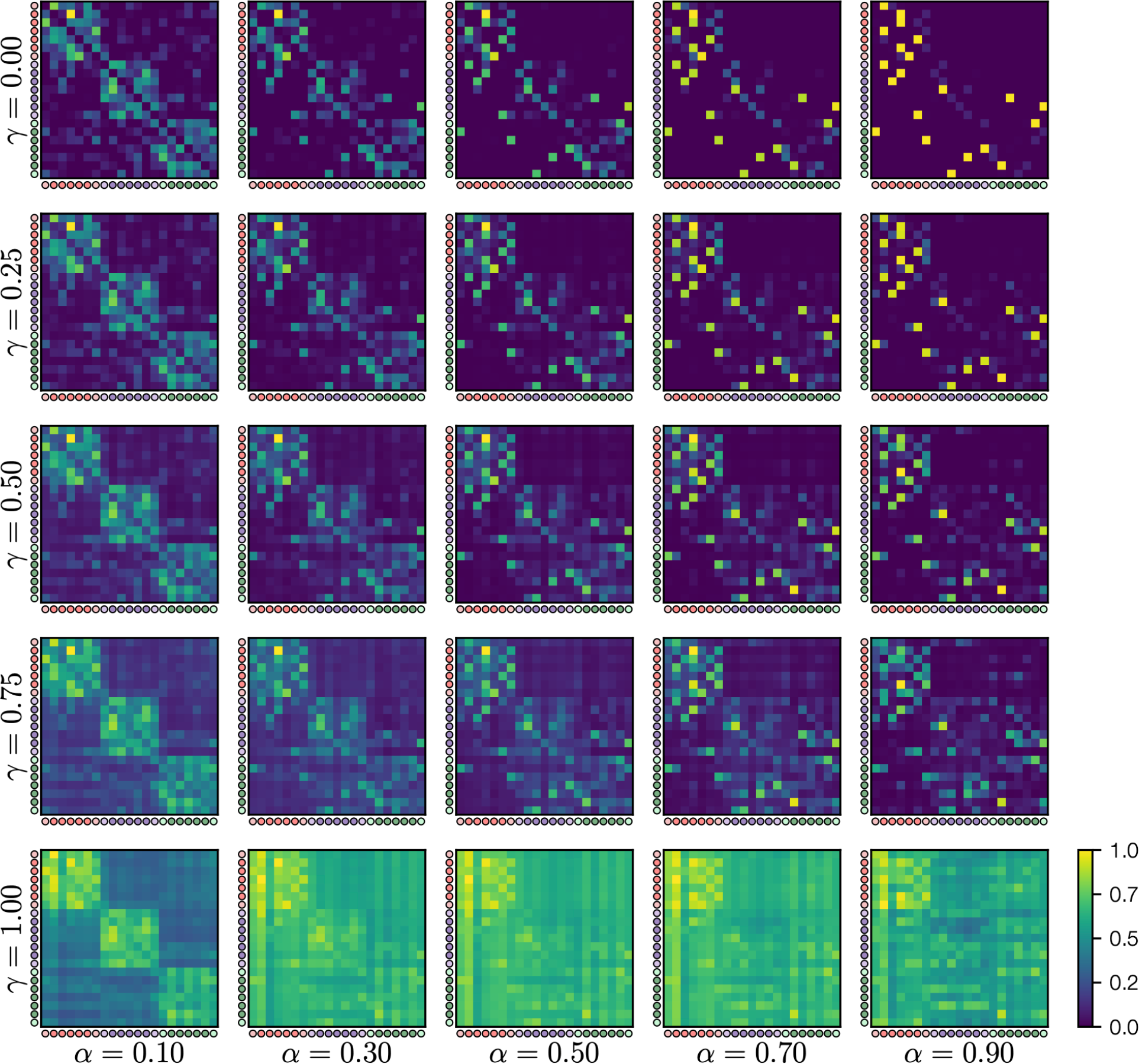
Successor representations after learning given different values of the *α* and *γ* parameters, after simulation of the object sequences viewed by an example participant. Within each successor representation, each row shows the expected count of future object presentations, discounted based on the discounting factor *γ*, for a given starting object. Circles indicate the object community (red, purple, green) and whether each object is a central node (dark) or boundary node (light). For display purposes, each matrix was divided by the maximum value in that matrix. We found that the clearest learning of the community structure appeared for relatively low values of *α* and relatively high values of *γ*.

Community structure after learning was most apparent when *α* was low, suggesting that gradual learning is important for detection of the underlying community structure. Based on these results, and following prior work (Stachenfeld et al., 2017), we fixed *α* = 0.1. The learned SR also varied based on the value of *γ* (Fig. 3). With *γ* = 0, only individual transitions are learned; as a result, indirect associations like those between the boundary items in a community are not reflected in the SR after learning. In contrast, with higher values of *γ*, both central and boundary items in the same community are associated with one another (Fig. 3).

We compared the SR model of object associations with a model that assumed that, after community structure learning, objects become associated with an abstract, reduced-dimensionality representation of the community structure. Abstract representations, which discard details of individual items to focus on common structure, have been theorized to guide decision making in a range of tasks, including associative inference, spatial navigation, and category learning (Mack et al., 2017; Morton et al., 2020; Stachenfeld et al., 2017; Varga et al.,n.d.). Here, we assumed that an abstract representation of the temporal community structure would focus on the community membership of each item, while discarding other details, such as whether two objects were directly or indirectly connected. We modeled the influence of abstract community representations on decision making by assuming that objects within the same community were associated (with strength 1) in an association matrix *W* and objects in different communities were not associated (strength 0). In this model, boundary items in the same community were just as strongly associated with one another as they were to central items, and boundary items in different communities were not associated at all (Fig. 5c).

We simulated the inference task based on the association strengths between the cue object *i* and the two choice objects *j* and *k* on each trial. The probability of choosing an object *j* was based on a Softmax choice rule, according to

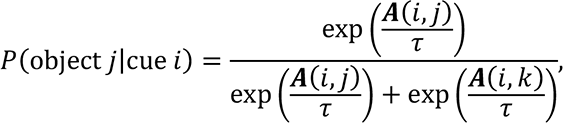

where *A* = *M* for the SR model, *A* = *W* for the community model, and *τ* is a temperature parameter. The *τ* parameter controls whether the object with highest association strength is selected with high probability (low *τ*) or whether responses are made more probabalistically, with both objects being selected equally often at high *τ*. For each model, we calculated the overall log-likelihood of the inference data by summing the log-transformed response probabilities across all trials.

We also considered a hybrid model where both SR and community strength matrices could contribute to inference decisions. A weight parameter *W*_*n*_, where *n* indicates the trial type, determined how strongly each matrix was weighted in determining inference responses:

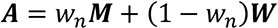

We hypothesized that the strength matrices might be differentially recruited on different trial types. For example, one might first retrieve memories of temporal associations (stored in the SR matrix), and only retrieve abstract community associations if no strong temporal associations are found in memory. In this case, abstract community associations would be most important on the boundary 2-away trials, wherein the cue was not directly associated with either of the object choices. To allow for such a possibility, in our model fits, the *w* parameter was estimated for each trial type separately.

Models were implemented in the publicly available Tesser package (Morton et al., 2021) using Python 3.7.6 with Numpy 1.18.1 (Harris et al., 2020) and Cython 0.29.23 (Behnel et al., 2010). We estimated model parameters using a differential evolution parameter search (Storn, 2008; Storn & Price, 1997) implemented in Scipy 1.4.1 (Virtanen et al., 2020), to estimate parameters that maximized the likelihood of the observed inference responses. The parameter search used the *best1bin* strategy (absolute tolerance=0, relative tolerance=0.01, mutation constant=[0.5, 1], recombination constant=0.7) with a population of 15 for each parameter being searched and Latin Hypercube initialization. After the initial differential evolution search, the best-fitting parameters were refined using the L-BFGS-B method (Byrd et al., 1995). Each search was run 10 times with different random starting parameters and the best-fitting parameters were selected across the different searches. Best-fitting parameters were then used to determine the fitted probability for each trial to select the within-community object. The temporal bias for each trial was calculated as 2(*P*(within) − 0.5) to obtain a measure where 1 indicates complete temporal bias toward inferring common properties for objects in the same community, 0 indicates no temporal bias, and -1 indicates a bias toward inferring common properties for objects in different communities.

Finally, we compared the SR, community, and hybrid models based on their log-likelihood and the number of free parameters in each model, using AIC (Akaike, 1998). AIC prefers models with higher log-likelihood and fewer free parameters. We also calculated AIC weights, which estimate the conditional probability of each model in a set being the best model, in terms of minimizing Kullback-Leibler discrepancy (Wagenmakers & Farrell, 2004).

### fMRI data collection

Imaging data were collected with a 3T Siemens Skyra at the Biomedical Imaging Center at The University of Texas at Austin. There were a total of six functional scans acquired using a T2*- weighted multiband-accelerated echo-planar imaging (EPI) pulse sequence (TR: 2000ms, TE: 30ms, flip angle: 73°, FOV: 220mm, 75 slices, 1.7x1.7x1.7 mm voxels, multiband factor: 3, GRAPPA factor: 2, phase partial Fourier: 7/8). A single-band reference (SBRef) image was also collected at the start of each EPI scan to aid in alignment to anatomical scans. A field map (TR = 628ms, TE = 5ms/7.46ms, flip angle = 5°, 64 slices, 1.7×1.7×2mm voxels) was collected prior to the first functional scan to correct for distortions in the magnetic field in functional scans 1–3. A second field map was collected after the break and used to correct for distortions in functional scans 4–6. Two oblique coronal T2-weighted (T2w) structural images were acquired perpendicular to the main axis of the hippocampus (TR: 13150ms, TE: 82 ms, flip angle: 150°, 60 slices, 0.4 x 0.4 mm in-plane resolution, 1.5 mm through-plane resolution) to facilitate segmentation of hippocampal anatomy. These images were co-registered and averaged to generate a mean coronal image for each participant. A T1-weighted 3D MPRAGE volume was collected (TR: 1900ms, TE: 2.43ms, flip angle: 9°, FOV: 256 mm, 192 slices, 1x1x1mm voxels) to facilitate alignment and normalization of the functional data.

### fMRI data preprocessing

Imaging data were converted to Brain Imaging Data Structure (BIDS) format 1.4.1 (K. J. Gorgolewski et al., 2016) using Heudiconv 0.9.0 (Yaroslav et al., 2020) and custom Python 3.7.6 scripts (Morton et al., 2021). Preprocessing of imaging data was performed using fMRIPrep 20.2.1 (O. Esteban et al., 2018; O. and M. Esteban, 2020), which is based on Nipype 1.5.1 (O. Esteban et al., 2020; K. Gorgolewski et al., 2011).

### Anatomical data preprocessing

The T1w image was corrected for intensity non-uniformity (INU) with *N4BiasFieldCorrection* (Tustison et al., 2010), distributed with ANTs 2.3.3 (Avants et al., 2008), and used as T1w- reference throughout the workflow. The T1w-reference was then skull-stripped with a Nipype implementation of the *antsBrainExtraction.sh* workflow (from ANTs), using OASIS30ANTs as target template. Brain tissue segmentation of cerebrospinal fluid (CSF), white-matter (WM) and gray-matter (GM) was performed on the brain-extracted T1w using *fast*, distributed with FSL 5.0.9 (Zhang et al., 2001). Brain surfaces were reconstructed using *recon-all*, distributed with FreeSurfer 6.0.1 (Dale et al., 1999), and the brain mask estimated previously was refined with a custom variation of the method to reconcile ANTs-derived and FreeSurfer-derived segmentations of the cortical gray-matter of Mindboggle (Klein et al., 2017). Volume-based spatial normalization to a standard space (MNI152NLin2009cAsym) was performed through nonlinear registration with *antsRegistration* (ANTs 2.3.3), using brain-extracted versions of both the T1w reference and the T1w template. The following template was used for spatial normalization: ICBM 152 Nonlinear Asymmetrical template version 2009c (Fonov et al., 2009). Hippocampus was automatically segmented based on both the T1w and averaged T2w scans using the *segmentHA_T2* program from FreeSurfer 7.1.1. (Iglesias et al., 2015; Saygin et al., 2017).

### Functional data preprocessing

For each of the six BOLD runs per subject, the following preprocessing was performed. First, a reference volume and its skull-stripped version were generated from the SBRef scans. A B0- nonuniformity map (or field map) was estimated based on a phase-difference map calculated with a dual-echo GRE (gradient-recall echo) sequence, processed with a custom workflow of SDCFlows inspired by the *epidewarp.fsl* script and further improvements in HCP Pipelines (Glasser et al., 2013). The fieldmap was then co-registered to the target EPI reference run and converted to a displacements field map (amenable to registration tools such as ANTs) with FSL’s *fugue* and other SDCflows tools. Based on the estimated susceptibility distortion, a corrected EPI reference was calculated for a more accurate co-registration with the anatomical reference.

The BOLD reference was then co-registered to the T1w reference using *bbregister* (FreeSurfer), which implements boundary-based registration (Greve & Fischl, 2009). Co-registration was configured with six degrees of freedom. Head-motion parameters with respect to the BOLD reference (transformation matrices, and six corresponding rotation and translation parameters) are estimated before any spatiotemporal filtering using FSL’s *mcflirt* (Jenkinson, 2002). BOLD runs were slice-time corrected using *3dTshift* from AFNI 20160207 (Cox & Hyde, 1997). The BOLD time-series were resampled into the standard MNI152NLin2009cAsym space with native functional resolution (1.7 mm isometric).

In addition to head-motion estimates calculated during motion correction, confounding time-series were calculated based on the preprocessed BOLD: framewise displacement (FD), DVARS and average signals within white matter and cerebrospinal fluid. FD was computed as the relative root mean square displacement between affines (Jenkinson et al., 2002). The confound time series derived from head motion estimates and global signals were expanded with the inclusion of temporal derivatives for each. All resamplings were performed with a single interpolation step by composing all the pertinent transformations (i.e., head-motion transform matrices, susceptibility distortion correction, and co-registrations to anatomical and output spaces). Resamplings were performed using *antsApplyTransforms* (ANTs), configured with Lanczos interpolation to minimize the smoothing effects of other kernels (Lanczos, 1964). Many internal operations of fMRIPrep use Nilearn 0.6.2 (Abraham et al., 2014), mostly within the functional processing workflow. For more details of the pipeline, see the section corresponding to workflows in fMRIPrep’s documentation.

### Regions of interest

Based on prior work examining the neural representation of temporal community structure, we identified hippocampus, inferior frontal gyrus (IFG), and medial prefrontal cortex (mPFC) as *a priori* regions of interest (ROIs) (Schapiro et al., 2013, 2016). We defined a hippocampal ROI based on the automatic FreeSurfer segmentation of the T1w and T2w anatomical scans for each participant. Subject hippocampal head, body, and tail ROIs were transformed to the MNI152NLin2009cAsym template space using *antsRegistration* with MultiLabel interpolation, pooled to generate right and left hippocampus ROIs, and averaged across participants to create probabilistic masks. We then thresholded these masks at 0.25 to obtain right and left hippocampal ROIs in template space.

We defined IFG ROIs based on FreeSurfer segmentation of the T1w anatomical scans for each participant, including the pars opercularis, pars triangularis, and pars orbitalis labels in the ROI for each hemisphere. As for the hippocampal ROIs, individual subject ROIs were transformed to template space and averaged to create right and left probabilistic masks. The masks were then thresholded at 0.1. We followed a similar procedure to generate a gray matter ROI that included all cortical and subcortical regions of gray matter, thresholded at 0.1. An mPFC ROI was manually drawn on the MNI ICBM 152 Non-Linear 6th Generation Asymmetric (MNI152NLin6Asym) template based on published definitions of anatomical mPFC (Öngür et al., 2003; Price & Drevets, 2009). The mPFC ROI in MNI152NLin6Asym space was then transformed to our main template space (MNI152NLin2009cAsym) using *antsRegistractionSyN.sh* and *antsApplyTransforms* from ANTS 2.3.5 (Avants et al., 2010) and intersected with the gray matter mask to create the final ROI.

### Searchlight analyses

We used searchlight analyses to examine whether temporal community structure is represented in the brain during the scanned part of the structure learning task. We focused specifically on the scrambled blocks, in which objects were presented in random order, to control for temporal autocorrelation and avoid systematic effects of object presentation order. This approach allowed us to determine whether the representations of individual objects had been affected by exposure to the temporal community structure. First, we used the SUSAN tool from FSL 6.0.4 (Smith & Brady, 1997) to smooth the preprocessed BOLD images in template space, with a 4 mm FWHM smoothing kernel. We then used a general linear model to estimate the pattern of activation that was elicited in response to each object during the scrambled blocks in each run.

We used Nilearn 0.7.1 and custom Python 3.7.0 code (Morton et al., 2021) to estimate object patterns using the least squares–separate (LS-S) approach (Mumford et al., 2012).

Object presentations (each modeled with 1 s duration) were convolved with a canonical double-Gamma hemodynamic response function to generate the predicted response time series for each object, separately for the scrambled and structured blocks. The activation of each object in the scrambled blocks was estimated using a separate model; modeled responses to other objects during the scrambled blocks were summed to create a single separate regressor. Each model also included regressors for each object presented during the structured blocks as regressors of no interest. Models also included 16 nuisance regressors generated by fMRIprep, consisting of 6 motion parameters and their temporal derivatives, and white matter and cerebrospinal fluid average signals and their temporal derivatives. Finally, each model included cosine basis functions to model slow fluctuations at less than 128 Hz. The LS-S modeling resulted in a beta series estimate of the activation at each voxel for each object in each run during the scrambled blocks.

We then used a searchlight analysis implemented in Brainiak 0.11.0 (Kumar et al., 2020) to test for neural representations of community structure. Separate searchlights were carried out for gray matter, left hippocampus, right hippocampus, left IFG, right IFG, and mPFC. In each searchlight sphere, we calculated all pairwise correlations between object patterns estimated based on the scrambled blocks in all runs. We then defined two bins for within-community and across-community pairings. The within-community comparisons included all pairs that were objects in the same community, were different objects, and were presented in different runs.

The across-community comparisons were the same but only included pairs of objects that were in different communities. We calculated the mean within- and across-community similarity and took the difference between them to obtain a measure of neural community structure. This neural community structure statistic was compared to a baseline determined using a permutation test. Objects were randomly permuted the same way for each run, such that object identity was preserved but community structure was broken, and the neural community structure statistic was calculated. This process was repeated 1000 times to obtain a null distribution for the neural community structure statistic. The same permutation order was used across all searchlight spheres to preserve spatial structure in the permuted data. For each sphere, the actual statistic was compared to the permutation distribution to obtain a z-score. The *randomise* tool from FSL 5.0.11 was used to implement a sign-flipping test with 5000 permutations to determine voxel-level significance across subjects.

We next examined whether neural community structure, which would reflect temporal learning, predicted non-temporal inference decisions about object grouping and object properties. We first tested whether neural community structure predicted performance on the grouping task, which involved using temporal knowledge to determine a spatial grouping of objects. To measure correlation between neural community structure and the grouping task, we first calculated the average Euclidean distance between items in the same community in the final object positions from the grouping task. These distances were z-scored across subjects and multiplied by -1 to obtain a measure of relative tendency to cluster same-community objects. We then used *randomise* to assess correlation between neural community structure representation and community clustering in the grouping task, comparing actual correlations to a permutation distribution based on randomly permuting grouping task scores. The grouping task searchlight allowed us to identify brain regions that may support inference about object similarity and generalization of temporal learning to a spatial task. Finally, to identify brain regions involved in inductive inference about object properties, we assessed correlation between neural community structure and temporal bias on the inference task. We calculated average temporal bias for each participant, collapsing across trial types, and z-scored the bias scores across subjects. We then tested whether there was correlation between neural community structure representation and temporal bias using *randomise*.

To perform cluster correction of the searchlight results within each ROI, we used AFNI 21.1.07 (Cox et al., 2017). We first estimated the residuals from a beta series model the same as the one used to estimate object patterns, except that all objects were included in a single model. We then used *3dFWHMx* to estimate parameters of the spatial autocorrelation function for each run. We averaged these parameters over runs and participants to obtain estimated autocorrelation parameters for each ROI. Finally, we used *3dClustSim* to estimate cluster extent thresholds to control the false-positive rate at 0.05, using three nearest neighbor clustering, one-sided thresholding, and a voxel-wise threshold of 0.01.

### Regression analysis

We used a Bayesian regression analysis to assess whether there is a relationship between neural community structure representations and temporal bias on the inference task. For each significant cluster from our searchlight analyses, we first dilated the cluster mask by one voxel. For each participant, we then extracted the betaseries pattern within the dilated cluster mask. We then calculated the mean pattern correlation for items in the same community and the mean pattern correlation for items in different communities, using the same similarity bins as used in the searchlight analysis. Finally, we subtracted across-community similarity from within-community similarity, to obtain a measure of neural community representation for each participant and cluster.

We then used a Bayesian regression to assess whether temporal bias on any of the test trial types were related to neural community representation across participants. We used Bambi 0.7.1 (Capretto et al., 2020) to define a linear model with weakly informative priors (Capretto et al., 2020). Regression coefficient priors were defined as

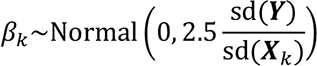

and the prior for the centered intercept was

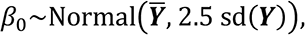

Where *Y* is the neural community representation for each participant, and the *X*_*k*_ predictors are the temporal bias scores for the central, boundary 1-away, and boundary 2-away trials.

We used PyMC 3.11.4 (Salvatier et al., 2016) to estimate the posterior distributions of the model parameters. Posterior distributions were estimated using the No U-Turn Sampler (NUTS) (Hoffman & Gelman, 2014) with 1000 tuning steps, target acceptance rate of 0.8, and 4 sampling chains with 2000 samples each. Convergence was assessed using bulk effective sample size and rank-normalized split potential scale reduction statistic Ȓ (Vehtari et al., 2019). For each model estimated and each parameter, Ȓ was less than 1.0001 and the bulk effective sample size was at least 4000. The posterior distribution of regression coefficients were estimated using the 95% high-density interval (HDI). For regression coefficients whose 95% did not include zero, we visualized the relationship by calculating the regression line based on the intercept and slopes for each posterior sample and then calculating the 95% HDI over that sample at each value of the predictor variable.

### Software and data availability

Code and Jupyter notebooks implementing the described analyses, including behavioral analysis, modeling, and representational similarity analysis, are publicly available (Morton et al., 2021). Behavioral and neuroimaging data will be made publicly available upon publication (Pudhiyidath et al., 2022).

## Results

### Performance during temporal structure learning

To assess whether participants consistently attended to the object sequences during temporal structure learning, we examined performance on the object orientation task. We calculated the sensitivity (measured using *d*′) of detecting objects that were rotated from their canonical orientation. Trials in which participants made no response, which were rare (relative frequency of no response: M=0.011, SE=0.002), were excluded from rotation detection analysis. Two runs (from different participants) were excluded due to errors in the recording of responses. The average detection sensitivity across all trials, including both the initial learning and scanned phases, was high (*d*′: M=4.023, SE=0.110). Detection sensitivity was above chance for each participant in each run (*p* < 0.001, permutation test).

To test whether fatigue affected performance during the structure learning task, we quantified whether performance varied between runs, either in initial learning phase that was not scanned, or during the scanned structure learning phase. We first used a one-way repeated measures ANOVA to test whether d’ varied across the five runs from the initial learning phase.

We observed a significant effect of run (*F*[4, 144] = 31.90, *p* < 0.001, η^2^_*p*_ = 0.47), with performance increasing during the first three runs of the initial learning phase. Follow-up *t*-tests of adjacent runs found significant increases in performance between runs 1 and 2 (change: M=0.688, SE=0.109, *t*(36) = 6.32, *p* < 0.001, Cohen’s *d* = 0.914, FDR-BH corrected) and runs 2 and 3 (change: M=0.403, SE=0.101, *t*(36) = 3.990, *p* < 0.001, *d* = 0.591, FDR-BH corrected).

There was a decrease in performance between runs 3 and 4 (change: M=-0.266, SE=0.082, *t*(36) = 3.24, *p* = 0.004, *d* = 0.411, FDR-BH corrected), but no change between runs 4 and 5 (*p* > 0.05, FDR-BH corrected). During the scanned structure learning, performance decreased but remained high (initial: M=4.208, SE=0.096; scanned: M=3.903, SE=0.066, *t*[36] = 3.60, *p* < 0.001, *d* = 0.950). There was no change in performance across the six scanned runs (*F*[5, 170] = 1.79, *p* = 0.117, η^2^_*p*_ = 0.050). Overall, these results suggest that attention was sustained throughout the task, after an initial period of acclimating to the task. However, because participants did vary in their overall performance (*d*′ range: 2.044 to 5.707), we included overall *d*′ as a covariate when examining individual differences in temporal knowledge expression, to control for individual differences in attention during temporal structure learning.

### Community structure predicts perception of temporal boundaries

Previous work found that exposure to temporal community structure affects the perception of event boundaries; when viewing sequences of objects, participants were more likely to indicate an event boundary after a transition between communities (Schapiro et al., 2013). We examined whether participants in our study exhibited similar event segmentation behavior during the parsing task. Specifically, we tested whether participants indicated an event boundary after transitions from one community to another, including only transitions that occurred after at least four consecutive steps into another community (Schapiro et al., 2013). We used a two-way repeated-measures ANOVA to examine whether parsing probability varied between object presentations that occurred at community transitions and other object presentations (Fig. 4a), and whether parsing depended on sequence type (Hamiltonian walk or structured random walk). We observed a significant main effect of parsing type (community transition vs. other; *F*[1,36] = 15.13, *p* < 0.001, η^2^_*p*_ = 0.296), no effect of sequence type (*F*[1,36] = 1.62, *p* = 0.212, η^2^_*p*_ = 0.043), and no interaction (*F*[1, 36] = 0.26, *p* = 0.614, η^2^_*p*_ = 0.007). These results are consistent with previous work (Schapiro et al., 2013) and demonstrate that participants perceived transitions between communities as event boundaries within the object sequence.

**Figure 4.**
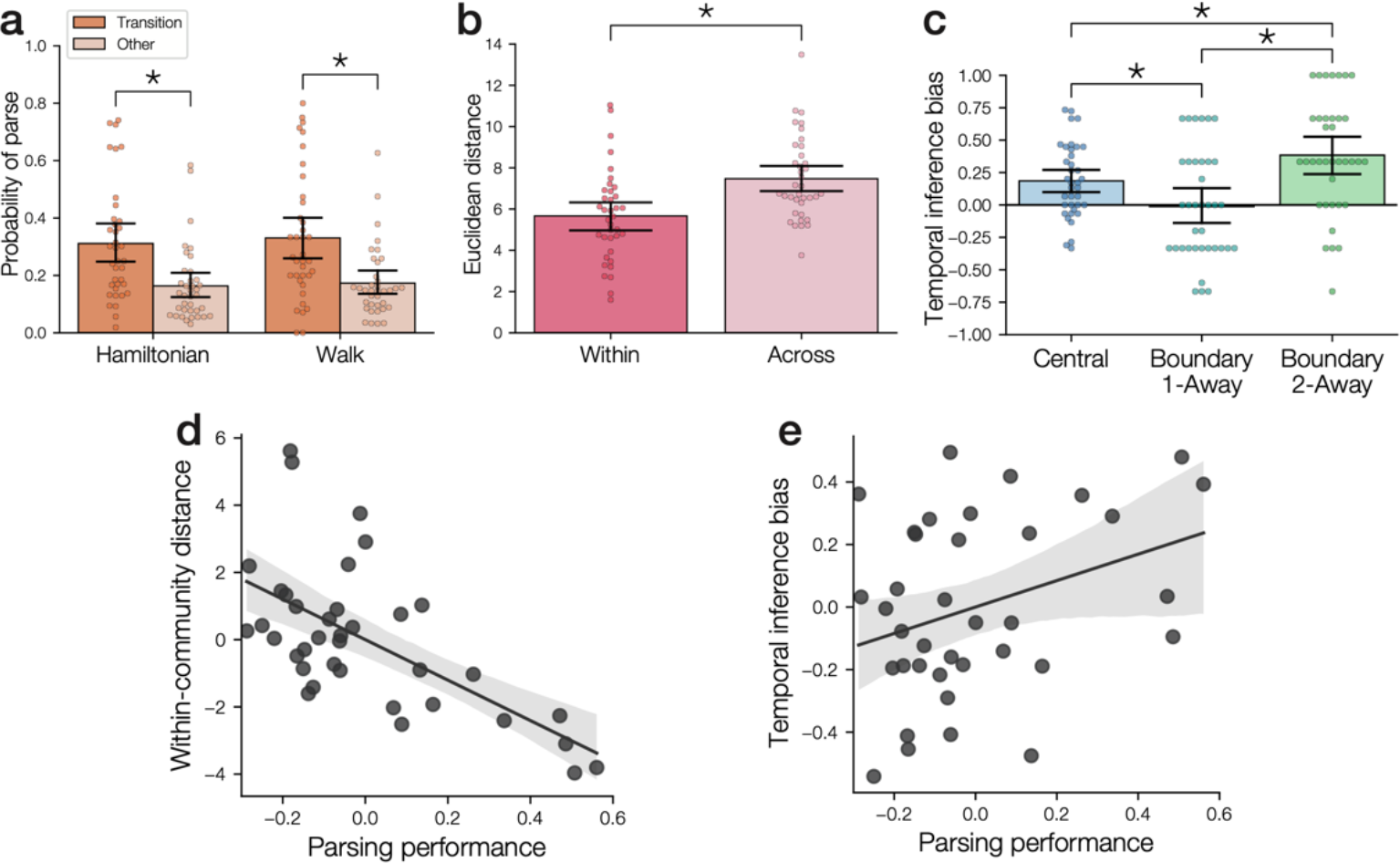
Behavioral measures of community structure learning. **(a)** Parsing probability during sequence viewing. Sequences were either Hamiltonian or random walks through the temporal community structure graph. Parsing occurred more frequently after transitions between communities compared to other transitions. Points indicate individual participants; error bars indicate bootstrap 95% confidence intervals. **(b)** In the grouping task, participants placed within-community objects nearer to one another than across-community objects. **(c)** Participants showed evidence of a bias toward inferring the same properties for objects from the same communities. Positive scores indicate bias toward inferring common properties of objects from the same community. **(d)** Parsing performance (i.e., event segmentation after community transitions relative to segmentation after other transitions) predicted smaller distances between objects in the same community on the grouping task. Both measures were residualized against rotation detection performance. Shaded area indicates 95% confidence intervals for the best-fitting linear regression. **(e)** Parsing performance predicted temporal bias on the inference task. Both measures were residualized against rotation detection performance. *: p<0.05.

### Subjective clustering of objects reflects community structure

We next examined whether statistical learning of the temporal associations between objects affected the subjective similarity of those objects, based on the spatial grouping task. While prior work demonstrated that temporal community structure affects segmentation of temporal sequences, it remains unclear whether knowledge about community structure is specific to temporal sequences of objects, or whether learning also shapes knowledge about the objects themselves. To test for an influence of community structure learning on the grouping task, we calculated the mean Euclidean distance between objects for within-community object pairs and across-community object pairs (Fig. 4b). We found that within-community objects were grouped closer together than across-community objects (*t*[36] = 4.56, *p* < 0.001, Cohen’s *d* = 0.854).

This result suggests that temporal community structure affects not only perception of temporal sequences, but also affects the subjective similarity of objects. Furthermore, our results demonstrate that object community membership, which was initially learned through exposure to temporal sequences, influenced behavior in a spatial task. These results thus suggest that temporal learning led to formation of an abstract representation of community that could influence performance in distinct tasks.

### Community structure biases inferences about object properties

We next tested whether temporal community structure affects reasoning about object properties. To assess how temporal knowledge affects inductive inference, participants were told a property of a cue object (i.e., the environment in which it is found; Fig. 2b) and asked which of two other objects shares that property. On each trial, one object was from the same community as the cue object, and one object was from a different community. Based on prior work (Pudhiyidath et al., 2019), we predicted that participants would be biased to select the object that shared the same community as the cue. While prior work focused on the overall tendency for community bias (Pudhiyidath et al., 2019), here we examined how bias varied depending on whether the cue item was directly or indirectly associated with each of the choice objects during learning. This analysis strategy allowed us to contrast the influence of different forms of temporal knowledge. Central trials required selecting between an object that was directly associated with the cue and an object that was indirectly associated with the cue. On these trials, inference decisions might be biased based on either knowledge of the direct association between the cue and the same-community object or knowledge that they shared the same community. On the boundary 1-away trials, both objects were directly associated with the cue object. We hypothesized that the direct association between the cue and the different- community object might reduce bias to select the same-community object. In contrast, on the boundary 2-away trials, neither object was directly associated with the cue object. These trials allowed us to examine the degree of same-community bias when object associations were indirect and therefore had to be inferred. We hypothesized that, if inference depended only on memory for direct associations between objects, then there would be no bias on the boundary2-away trials. In contrast, if inference depended on temporal community membership, then inference would be biased on the boundary 2-away trials.

First, we tested whether bias changed over time during the inference task; such a change might indicate that participants learned about the environment associated with each object during the inference task itself. A one-way repeated-measures ANOVA found that bias did not vary with trial bin (*F*[5,180] = 0.74, *p* = 0.594, η^2^_*p*_ = 0.020), confirming that temporal bias on the inference task reflected knowledge acquired during the structure learning phase rather than during the inference task itself. We next compared bias in inference responses for the different trial types. Consistent with our predictions, responses to central questions were significantly biased toward selection of within-community objects (*t*[36] = 3.97, *p* < 0.001, FDR-BH corrected, *d* = 0.652). In contrast, responses to boundary 1-away questions did not show bias toward one type of choice versus the other (*t*[36] = −0.18, *p* = 0.857, FDR-BH corrected, *d* = 0.030), suggesting that the direct association between the cue and the two choice items eliminated bias based on temporal community. Responses to boundary 2-away questions were significantly biased toward selecting the within-community objects (*t*[36] = 5.29, *p* < 0.001, FDR-BH corrected, *d* = 0.869), suggesting that participants did not rely solely on memory for direct associations, but were also biased by abstract knowledge about the temporal community structure.

Using a one-way repeated-measures ANOVA, we found a significant effect of question type (*F*[2,72] = 14.27, *p* < 0.001, η^2^_*p*_ = 0.284). Follow-up *t*-tests revealed significant differences between central and boundary 1-away trials (*t*[36] = 3.30, *p* = 0.003, *d* = 0.552), central and boundary 2-away trials (*t*[36] = 2.58, *p* = 0.015, *d* = 0.534), and boundary 1-away and boundary 2-away trials (*t*[36] = 4.74, *p* < 0.001, *d* = 0.920). These results suggest that the relationship between the objects has an important impact on inference of object properties. Furthermore, the bias toward selecting within-community objects becomes stronger when there was no direct connection between the cue object and the object whose properties are being inferred. These results suggest that statistical learning of community structure biases decisions more when participants do not have direct experience upon which to draw.

### Perception of event boundaries correlates with object clustering and inference

We observed that participants perceived a change in the object sequences when there was a shift from one community to another (Fig. 4a), suggesting that participants were sensitive to the community structure when segmenting events during the parsing task. We hypothesized that participants’ ability to parse the communities would predict individual differences in the two tasks that required generalization of knowledge to new task contexts—grouping and inductive inference. For each participant, we calculated the difference in parsing probability between community transitions and other object presentations to obtain an overall measure of parsing performance. We then tested whether parsing behavior correlated with spatial community clustering in the grouping task and temporal bias on the inference task (averaged over trial type). In each regression analysis, we used partial correlation to control for d’ during the rotation detection task as a measure of how well each participant attended to the object sequences during structure learning.

We observed a significant correlation between parsing performance and within-community distance in the grouping task (Fig. 4d; *r* = −0.628, *p* < 0.001) after controlling for rotation detection performance. There was no correlation between parsing performance and across-community distance (*r* = 0.065, *p* = 0.705) after controlling for rotation detection performance. This result is consistent with our observation that across-community distance varied substantially due to individual differences in the placement of groups within the response grid (see Methods). As across-community distance was not related to temporal knowledge, we focus on within-community distance for subsequent analysis of the grouping task. We also observed a significant correlation between parsing performance and temporal bias on the inference task, after controlling for rotation detection performance (Fig. 4e; *r* = 0.336, *p* = 0.045). These results suggest that the participants who were most knowledgeable about the boundaries of the communities were more likely to group communities together spatially after learning and more likely to use their knowledge of the temporal communities to make decisions about shared object properties. Finally, we examined whether object clustering in the grouping task was related to temporal bias on the inference task, after controlling for rotation detection performance. We did not find evidence of a relationship of temporal bias with within-community distance (*r* = −0.185, *p* = 0.282), suggesting that the grouping and inference tasks might reflect somewhat distinct cognitive processes.

### Inference decisions depend on both transition statistics and community structure

To better understand how temporal community structure learning biases inference decisions about object properties, we developed a model of statistical learning and inference in our task.

We found previously that inference behavior may depend on both knowledge of direct associations between objects and knowledge of temporal community membership. To examine these distinct influences on behavior, we compared models of temporal knowledge at different levels of abstraction. We first modeled statistical learning of object associations based on the successor representation (SR), which efficiently stores information about experienced sequences of states (Dayan, 1993; Momennejad, 2020; Momennejad et al., 2017; Stachenfeld et al., 2017). Given some current state (here, a currently presented object), the SR encodes a temporally discounted estimate of how frequently other states (i.e., other objects) will be visited in the future. Objects that are likely to be visited sooner will be weighted more strongly, but objects that are indirectly connected will also be represented as potential future states. We simulated learning of an SR using temporal difference learning (Momennejad, 2020; Stachenfeld et al., 2017). For each participant, the SR model was presented with the object sequence viewed by that participant. For each inference test trial, we took the current “state” as the cue object and calculated the predicted temporally discounted frequency at which each of the two choice objects would be visited from that cue object state (Fig. 5a). We assumed that the object with the higher predicted frequency was more likely to be selected.

**Figure 5.**
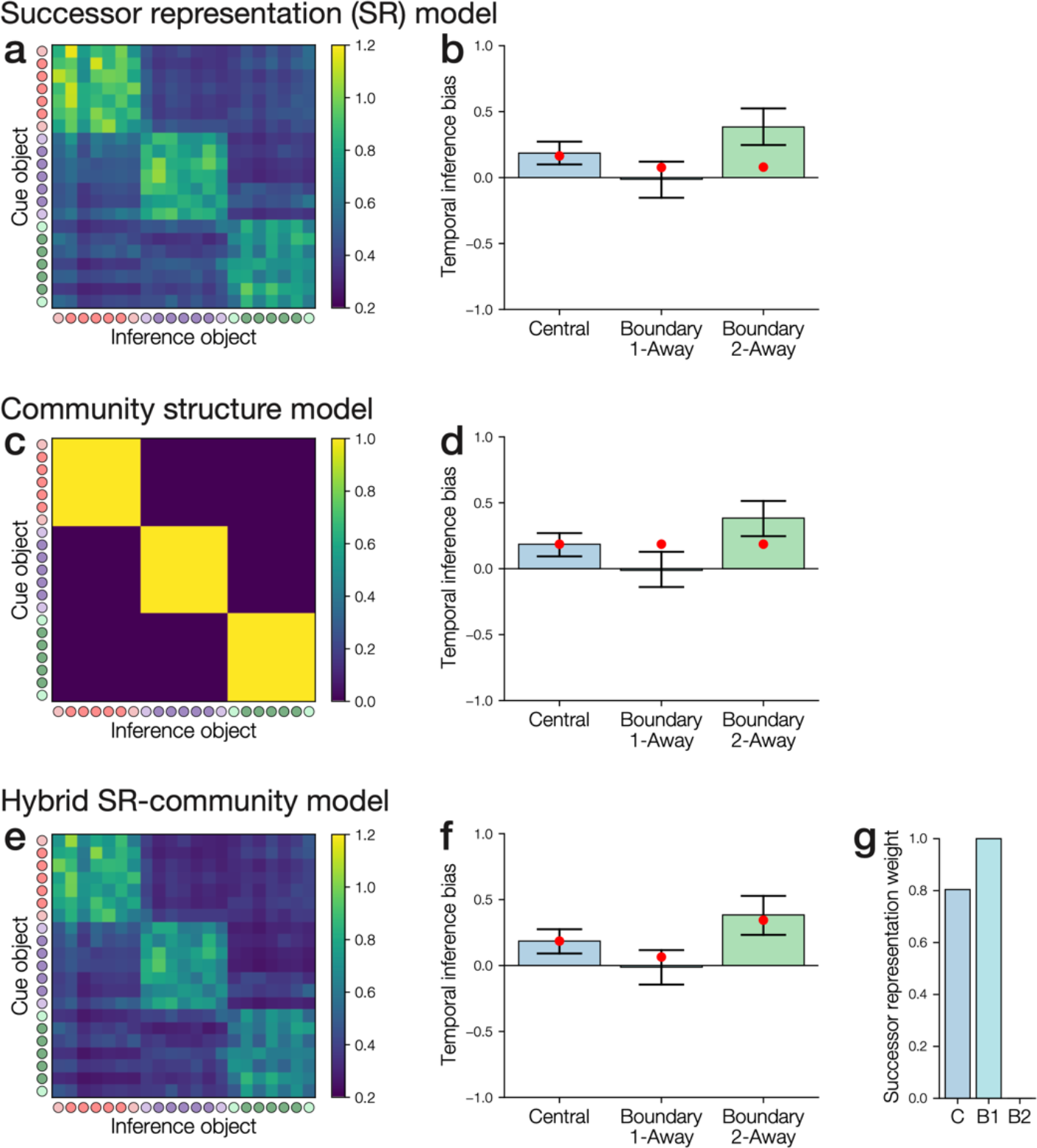
Models of inductive inference task performance based on learned similarity of object pairs. **(a–b)** Successor representation (SR) matrix and model fit to inference behavior. **(a)** SR matrix after learning for one example participant (the same participant as in Fig. 3) based on the object sequences that they observed, with a learning rate of 0.1 and best-fitting discounting factor of 1.0. The matrix shows the expected temporally discounted frequency of visiting each inference object after viewing the cue object. Circles indicate the object community (red, purple, green) and whether each object is a central node (dark) or boundary node (light). **(b)** Fit of the SR model to average temporal bias for each inference test type. Bars show observed performance (with bootstrap 95% confidence intervals); red dots show model estimates. The model incorrectly predicts that temporal bias should be low on the boundary 2-away trials. **(c–d)** Within-community similarity model. **(c)** Model association matrix. Objects in the same community have a similarity of 1 and objects in different communities have a similarity of 0. **(d)** The model incorrectly predicts that there will be equal bias for each trial type. **(e–g)** Hybrid SR-community model. The hybrid model simulates inference as being influenced by a weighted combination of SR and within-community similarity. **(e)** SR matrix for the same example participant as in **(a)**, with the best-fitting discounting factor of 0.974. **(f)** The hybrid model correctly fits the differences in temporal bias between the trial types. **(g)** Based on the model fit, the central and boundary 1-away inference trials weight the SR more heavily, while the boundary 2-away questions weight only the within-community similarity matrix. C: central; B1: boundary 1-away; B2: boundary 2- away.

While the SR reflects detailed information about transition probabilities, we hypothesized that participants might also learn an abstract representation of the objects that purely reflects their community membership (Fig. 5b). We assumed that, in this case, the cue would support inference of shared properties with any objects in the same community, but not objects from other communities. This model thus assumes that inference decisions will not depend on whether objects were directly associated during learning, but instead only based on whether they were in the same community as the cue object.

We first examined whether either the SR model or the community membership model could account for the temporal bias observed for the central, boundary 1-away, and boundary 2- away trials. We found that the SR model with best-fitting parameters (gamma=1.000, tau=1.337) provided a reasonable fit to the central and boundary 1-away trials, but incorrectly predicted that boundary 2-away questions should exhibit less temporal bias (Fig. 5a). In contrast, the community membership model with best-fitting parameters (tau=2.655) incorrectly predicted that all inference trials should exhibit the same amount of temporal bias, as the same-community object was always favored regardless of how closely connected it or the other object were to the cue object (Fig. 5b).

Neither the SR or community membership models could completely account for the pattern of bias on the inference task. This finding is consistent with our behavioral results, which suggest that the boundary 1-away and boundary 2-away trials may rely on distinct forms of temporal knowledge. Specifically, there was no bias when both choices were directly associated with the cue (boundary 1-away), suggesting that direct associations influence inference behavior. In contrast, we observed strong bias when neither choice as directly associated with the cue (boundary 2-away), suggesting that participants also learn about temporal community membership. We therefore next examined whether a hybrid model using both SR and community membership could fit performance on the inference task. Inference decisions were modeled as being based on a weighted combination of SR and community membership. We estimated the relative weight of the components separately for each trial type. The hybrid model obtained a good fit to the temporal bias observed for each of the inference trial types (Fig. 5c). We compared the fits for the SR, community, and hybrid models to inference test performance, while controlling for model complexity. We found that the hybrid model performed the best overall (wAIC=0.989, AIC=2697.630), followed by the community model (wAIC=0.011, AIC=2706.568) and the SR model (wAIC=0.00006, AIC=2717.155). Examining the best-fitting parameters of the hybrid model, we found that, while the central trials and boundary 1-away trials mostly relied on the SR, boundary 2-away trials exclusively weighted community membership (Fig. 5c; the other best-fitting parameters were: *γ* = 0.974, *τ* = 1.392). In both the central and boundary 1-away trials, responses may be informed by direct experience, as the cue object was directly connected to at least one of the choice objects. In contrast, in boundary 2-away trials, participants must choose between two objects that were never presented adjacent to the cue object (except rarely, during the scrambled blocks of the scanning task; see Fig. 2b). On these trials, we found evidence that participants instead make inference decisions based on the more abstract knowledge of the temporal community structure, resulting in a stronger temporal bias.

Our modeling results suggest that that the boundary 1-away and boundary 2-away trials are particularly distinct, relying on an SR-like representation and a community-like representation, respectively. Boundary 1-away responses may depend on an SR-based mechanism, while boundary 2-away responses may depend on a more abstract representation of community structure.

### Anterior hippocampus represents temporal community structure

We next examined whether exposure to temporal community structure affects the neural representation of individual objects. A previous study observed that, after exposure to temporal community structure, Hamiltonian sequences of objects elicited activation patterns in IFG (Schapiro et al., 2013) and hippocampus (Schapiro et al., 2016) that were more similar for objects in the same community. The authors also observed that activation in IFG increased during a series of transitions within a given community, suggesting that the order in which the objects are presented may affect brain activation in response to object presentations. In the present study, we measured activation patterns of each object in the scrambled blocks of the scanning task (Fig. 2a), allowing us to quantify object representations in the absence of any predictable ordering. We tested whether objects in the same community were represented more similarly than objects in different communities, only comparing patterns across different scanning runs to control for pattern correlations within runs (Mumford et al., 2014). We used separate searchlight analyses to identify regions representing community structure, looking within a priori ROIs of right and left hippocampus, right and left IFG, and mPFC. Finally, a broader searchlight analysis included all cortical and subcortical grey matter.

We found that activation patterns in right anterior hippocampus were more correlated for objects in the same community than objects in different communities (Fig. 6; *p* < 0.05, cluster-corrected within the volume of the searchlight). The other searchlights did not reveal any significant clusters (*p* > 0.05, cluster-corrected). This result extends prior work showing evidence of temporal community structure representation in right anterior hippocampus (Schapiro et al., 2016) by demonstrating that object representations in hippocampus are sensitive to temporal community structure even when object presentation order is scrambled. In contrast to prior work (Schapiro et al., 2013), we did not observe any significant clusters in IFG. This difference may be due to the lack of a predictable order to the presented objects in the scrambled blocks, in contrast to the prior study in which objects were presented in an order taken from a Hamiltonian path through the temporal community structure graph (Schapiro et al., 2013). This prior study found that IFG activity increased over multiple presentations within a community (Schapiro et al., 2013), which would not generally happen in the present study due to the random order in which objects were presented.

**Figure 6.**
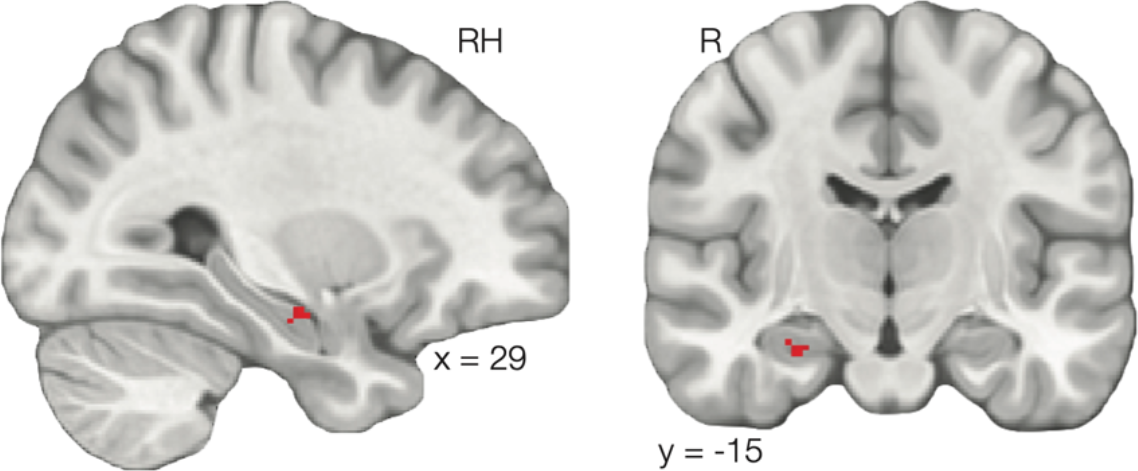
Right anterior hippocampus similarity is greater for objects in the same community compared to objects in different communities.

### Pattern similarity in perirhinal cortex predicts subjective object similarity

We next examined whether neural pattern similarity predicted individual differences in the subjective similarity of objects in the grouping task. While prior work found that neural representations reflect temporal community structure (Schapiro et al., 2013, 2016), it remains unclear whether these representations are related to behavior. To characterize the functional significance of neural community representations, examined whether the similarity contrast (within- minus across-community similarity) for each participant was correlated with within-community similarity from the grouping task. The whole-brain searchlight revealed that left perirhinal cortex was significantly correlated with within-community distance in the grouping task (Fig. 7a). Searchlights within hippocampus, IFG, and mPFC did not reveal any significant clusters (*p* > 0.05, cluster-corrected). These results suggest that, in participants who grouped together objects in the same community more closely during the grouping task, object representations in perirhinal cortex reflected the temporal community structure. We propose that learning of temporal community structure may shape high-level object representations in perirhinal cortex, which may then affect the subjective similarity of the objects, leading to tighter clustering of within-community objects in the grouping task.

**Figure 7.**
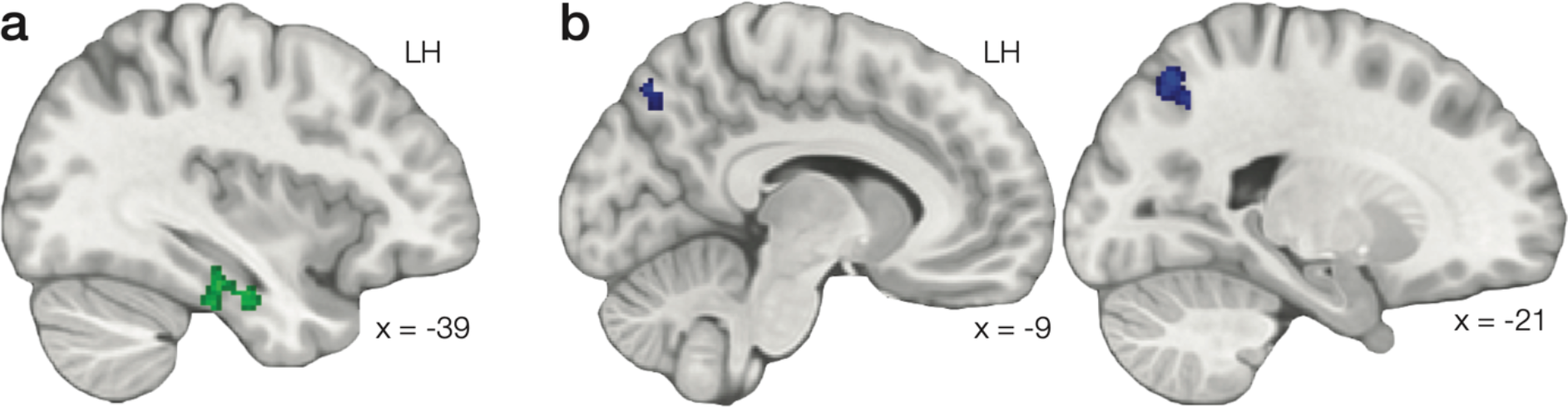
(a) A greater difference between within- and across-community similarity in left perirhinal cortex predicted smaller distance between same-community objects in the grouping task. **(b)** Left precuneus showed a greater difference between within- and across-community similarity for participants that were more biased by community structure on the inference task.

### Pattern similarity in precuneus and hippocampus predicts inference

We next examined whether neural pattern similarity is related to individual differences in inference. We tested whether within-community minus across-community similarity was related to temporal bias on the inference task, averaged over all trials for each participant. The whole-brain searchlight revealed a region in left precuneus where neural representation of object community structure predicted increased bias on the inference task (Fig. 7b; *p* < 0.05, cluster corrected). Searchlights within hippocampus and IFG did not reveal any significant clusters (*p* > 0.05, cluster corrected). Our results suggest that presentation of individual objects may result in activation of community-specific representations in precuneus; these representations might subsequently influence decision-making when participants are asked to infer which objects come from the same environment. We next used a Bayesian regression analysis to examine whether precuneus pattern similarity was uniquely related to any specific inference test trial type. This analysis allowed us to estimate the strength of relationship of precuneus pattern similarity to each test trial type. We found evidence of a positive relationship between precuneus pattern similarity difference and temporal bias on the boundary 1-away trials (Fig. 8a; slope 95% high-density interval [HDI]: [0.001, 0.032]). We did not find clear evidence of a relationship on the central (slope 95% HDI: [-0.001, 0.044]) or boundary 2-away trials (slope 95% HDI: [-0.021, 0.005]).

**Figure 8.**
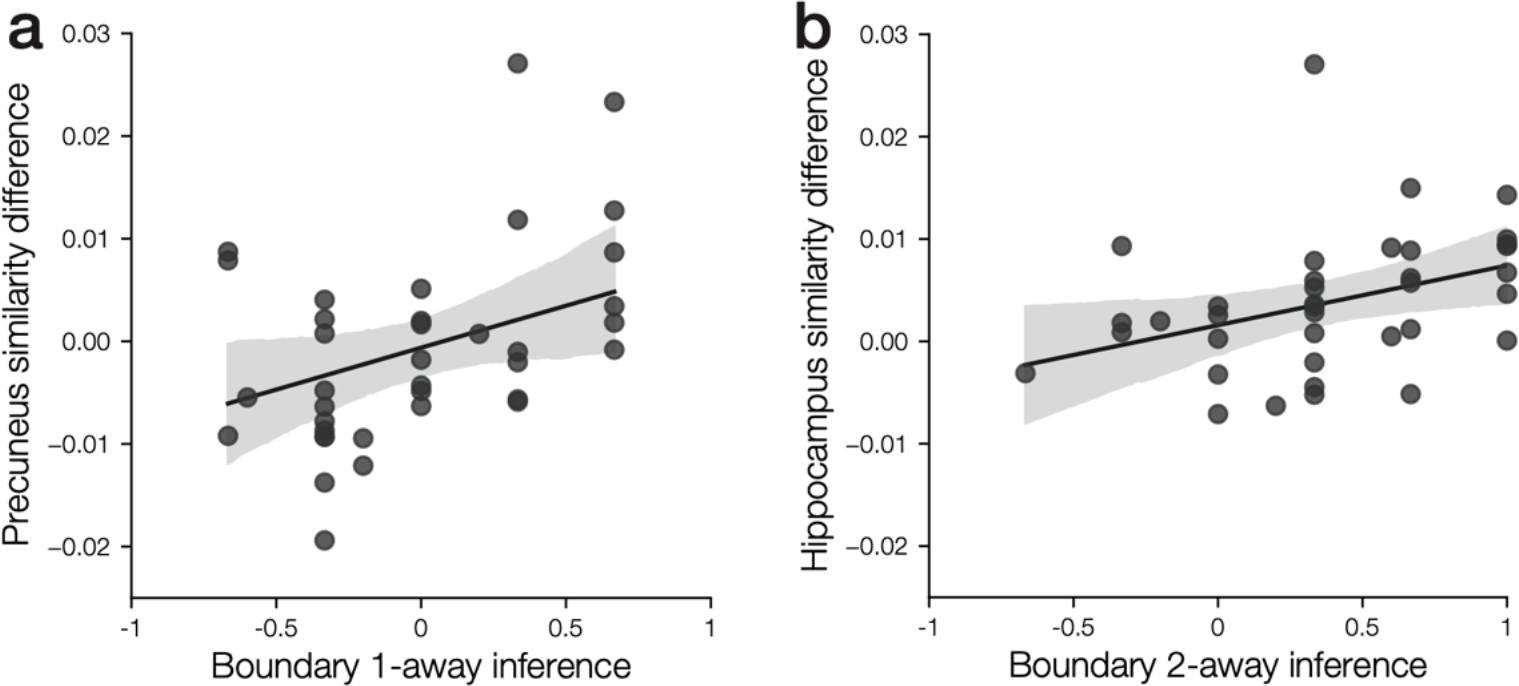
(a) Community structure in the left precuneus cluster was uniquely related to temporal bias on the boundary 1-away inference trials. **(b)** Community structure in the right anterior hippocampus cluster was uniquely related to temporal bias on the boundary 2-away inference trials.

While there were no significant clusters in hippocampus or perirhinal cortex in the inference temporal bias searchlight, our behavioral and modeling results suggest that the different inference tests may involve distinct mechanisms. Therefore, we used a Bayesian regression analysis to estimate the strength of the relationships between bias on the individual inference trial types and the hippocampal or perirhinal cortex community pattern similarity. We found evidence of a positive relationship between hippocampal pattern similarity differences and temporal bias on the boundary 2-away trials (Fig 8b; slope 95% HDI: [0.001, 0.022]). We did not find clear evidence of a relationship on the central (slope 95% HDI: [-0.028, 0.009]) or boundary 1-away (slope 95% HDI: [-0.010, 0.015]) trials. We did not find clear evidence of a relationship between perirhinal pattern similarity differences and temporal bias on the central (slope 95% HDI: [-0.002, 0.021]), boundary 1-away (slope 95% HDI: [-0.014, 0.003]), or boundary 2-away (slope 95% HDI: [-0.001, 0.013]) trials. Overall, our results suggest that pattern similarity in hippocampus and precuneus is related to effects of temporal community structures on inference decisions about object properties. Furthermore, these relationships may be selective, with precuneus related to performance on the boundary 1-away trials and hippocampus related to performance on the boundary 2-away trials.

## Discussion

Previous work found that exposure to temporal community structure biases inductive inference (Pudhiyidath et al., 2019), suggesting that knowledge of latent structure affects reasoning about object properties. However, the cognitive and neural mechanisms through which temporal statistical learning influences reasoning remained unclear. Here, we found evidence that temporal community structure affects both inference decisions about object similarity and inductive inference decisions about object properties. The fidelity of community structure representations in perirhinal cortex predicted the degree to which objects were grouped by community, suggesting that statistical learning may influence high-level object representations, which then bias similarity judgments. We found that temporal knowledge also biases inductive inference decisions about properties shared among different objects. Using computational modeling, we found that both detailed knowledge of temporal sequences and abstract knowledge of temporal community structure influence inductive inference decisions. These inductive inference decisions may be supported by hippocampus and precuneus, which represent the temporal community membership of objects more coherently for participants who exhibit stronger temporal bias during inductive inference.

We first examined whether temporal community structure affected perception of object similarity by asking participants to group the objects on a spatial grid. We found that objects that had been in the same community during learning were more likely to be grouped together. This result demonstrates that knowledge of latent community structure, which was originally learned through exposure to temporal sequences, influences behavior in a non-temporal spatial grouping task. The influence of temporal learning on object grouping may be driven by changes to object representations in perirhinal cortex. Participants who exhibited more coherent neural representations of temporal community in perirhinal cortex after temporal learning tended to group together objects in the same community. Perirhinal cortex is situated at the apex of the ventral visual processing stream (Miyashita, 2019; Suzuki & Amaral, 1994), and in contrast to earlier visual areas, object representations in perirhinal cortex reflect learned associations between objects (Naya et al., 2001; Sakai & Miyashita, 1991; Schapiro et al., 2012).

Furthermore, perirhinal cortex is necessary for reactivation of associated objects in earlier visual areas (Higuchi & Miyashita, 1996; Miyashita, 2019) and has been proposed to help support memory for associations between items (Mayes et al., 2004, 2007; Ranganath, 2010b; Yakovlev et al., 1998). Previous work found that object representations in perirhinal cortex reflect temporal proximity in a repeated sequence; objects nearer to one another had more similar representations after learning (Miyashita, 1988). Our results further suggest that object representations in perirhinal cortex are also sensitive to latent community structure. The formation of associations between items based on shared temporal context, as we observed here, has been proposed to be a key mechanism for learning about semantic relationships (Howard et al., 2011; Landauer & Dumais, 1997). Previous work suggests that neural representations in perirhinal cortex reflect semantic associations, which may help to disambiguate objects that are visually similar (Barense et al., 2010; Martin et al., 2018). Taken together with prior studies, our results suggest that learning of latent community structure may facilitate disambiguation of objects in perirhinal cortex based on their associated context. The representational differences among objects in different communities may then influence decisions about the subjective grouping of the objects.

In addition to influencing judgments of object similarity, temporal community structure biased inductive inference about object properties. This bias was greater for participants who exhibited consistent parsing at transitions between communities, suggesting that formation of temporal knowledge during sequence viewing caused bias based on temporal associations during inductive inference. Our results confirm prior findings that temporal community structure learning leads to biased inductive inference (Pudhiyidath et al., 2019). Critically, however, we found that the bias toward inferring shared properties among objects in the same community was strongest when participants had not learned any direct associations between the cue and either choice object (i.e., on boundary 2-away trials). In contrast, when both choice objects had been directly associated with the cue object (i.e., on boundary 1-away trials), there was no bias to select the object from the same community.

Using computational modeling, we found that this pattern of results could be explained if inductive inference decisions were based on a combination of knowledge about object transition probabilities, as encoded in an SR (Dayan, 1993; Momennejad et al., 2017; Stachenfeld et al., 2017) and an abstract low-dimensional representation of community membership. The SR has been proposed to support discovery of clusters (here, object communities) within states (here, the objects) (Momennejad, 2020; Stachenfeld et al., 2017), suggesting that an SR-like learning mechanism may support formation of both predictive representations of transition statistics and abstract representations of community membership. Our modeling results suggest that abstract community representations only influence inductive inference behavior when there are no direct associations between the cue and the objects being considered. This shift in emphasis may be adaptive, focusing on direct experience when possible, and only relying on inferred latent structure when direct experience is not available. Our results are consistent with theoretical work suggesting that the temporal features of experience support inference about latent causes (Gershman et al., 2017). Latent cause inference has been proposed to be a critical mechanism for organizing memory, as it may allow individual experiences to be connected to detailed schemas that can guide reasoning (Varga et al., n.d.).

Our modeling results are consistent with the hypothesis that inductive inference is influenced, in part, by predictive representations of temporal relationships between objects. An alternative model of inductive inference, however, could posit that object properties are instead generalized from a given cue object to objects that tend to preceed the cue object during learning. Such a mechanism would be consistent with temporal context models, which propose that presented items become associated with a contextual trace that represents recently presented items (Gershman et al., 2012; Howard & Kahana, 2002). Because the community structure graph in the present study was bidirectional, these different models of inference would likely give similar predictions for behavior in the inductive inference task. To help clarify further how inductive inference is influenced by temporal experience, a future study could present participants with object sequences following a directional graph. An inductive inference task could then be used to examine whether object properties are inferred to be shared with preceding objects, predicted upcoming objects, or both. Regardless of the directionality of the temporal associations that influence inference behavior, however, our results suggest that inductive inference behavior is sensitive to both the community membership of the objects and the temporal distance between the objects during learning.

We found evidence that inductive inference based on latent community structure is supported by neural representations of community structure in the hippocampus. The hippocampus has previously been implicated in supporting statistical learning (Covington et al., 2018; Schapiro et al., 2013; Schlichting et al., 2017), potentially by forming SR-like predictions of upcoming states (Stachenfeld et al., 2017). The hippocampus is also thought to support binding of items with their associated contexts (Ranganath, 2010a; Tort et al., 2009). Here, our data suggest that the hippocampus forms associations between objects and their latent community context. We reproduced prior results finding community representations in right anterior hippocampus (Schapiro 2016), and further found that this property persists when objects are not presented in a predictable order. Hippocampal representations of community membership are thus activated in response to individual objects rather than only appearing during structured temporal sequences drawn from the temporal community structure graph.

Hippocampus has also been proposed to integrate indirectly related items (Molitor et al., 2021; Schlichting et al., 2015); this mechanism of memory integration has been proposed to serve a critical role in deductive inference based on indirect associations (Bunsey & Eichenbaum, 1996; Molitor et al., 2021; Schlichting & Preston, 2015; Varga et al., n.d.). Consistent with the hypothesized role of hippocampus in supporting inference based on indirect associations, we found that the strength of temporal community structure representations in hippocampus was uniquely related to biased inference on the boundary 2-away trials, which required making decisions based on indirect associations. Integrative representations in hippocampus have previously been linked to deductive inference (i.e., given learned pairings of AB and BC items, A and C are also associated) (Molitor et al., 2021). Here, we find that hippocampal representations also predict inductive inference decisions, wherein a decision cannot be deduced from existing knowledge and must instead be extrapolated based on abstract knowledge of latent structure.

We found that neural representations of community structure in precuneus were also related to bias in the inductive inference task. The precuneus has previously been found to be involved in encoding and retrieving contextual associations (Frings et al., 2006; Sreekumar et al., 2018; Wagner et al., 2005) and representing latent structure (Baldassano et al., 2017, 2018; Vaidya et al., 2021), suggesting that it may be involved in representing the community associated with each object. Furthermore, precuneus is activated during decisions about whether two objects are often found in the same context (e.g., a washer and a laundry hamper) or not (e.g., a microscope and a toy horse), and activation is greater when the pair of objects is related (Livne & Bar, 2016). The precuneus may similarly be recruited during inductive inference to determine whether two objects are associated with the same community and thus might share some of the same properties. We also found initial evidence that the precuneus may be selectively predictive of inductive inference decisions when items were directly associated during learning. Our modeling results suggest that these inductive inference decisions were driven by detailed temporal knowledge. Taken together, the neural and modeling results suggest that precuneus may support inductive inference decisions based on detailed temporal knowledge of object associations. Presentation of a cue object during the inductive inference task might trigger reactivation of associated latent task structure in precuneus. Representations of latent structure might then bias inductive inference decisions to favor generalization to objects that are associated within that latent structure.

In conclusion, we found that learning about statistical temporal associations influences both inference decisions about object relationships and inductive inference decisions about shared object properties. Reasoning decisions that require inductive inference appear to reflect retrieval of both temporal associations between objects, which may be supported by precuneus, and abstract community structure, which may be supported by hippocampus. Inference about latent causes, such as shared associations and temporal community structure, has been proposed to be a critical organizational feature of memory (Gershman et al., 2017). Our results suggest that latent cause inference may be influenced by multiple features of experience that compete to determine memory-based reasoning behavior.

## Acknowledgements

The authors thank Susannah Cox, Jillian Perez, and Ata Karagoz for their assistance in collecting the data and for helpful discussions; and Jackson Liang for help with defining anatomical masks.

## Notes

https://doi.org/10.18112/openneuro.ds004094.v1.0.0

https://doi.org/10.5281/zenodo.4793426

### Competing Interest Statement

The authors have declared no competing interest.

### Summary of Updates

A new analysis has been added to demonstrate the effects of model parameters on learning of the successor representation (Fig. 3); more detail has been added about the parameter search methods; brain-behavior correlation analyses have been revised to use Bayesian methods; additional interpretation of model results has been added to the discussion.

